# Uncoating of COPII from ER exit site membranes precedes cargo accumulation and membrane fission

**DOI:** 10.1101/727107

**Authors:** Olga Shomron, Inbar Nevo-Yassaf, Tamar Aviad, Yakey Yaffe, Eitan Erez Zahavi, Anna Dukhovny, Eran Perlson, Ilya Brodsky, Adva Yeheskel, Metsada Pasmanik-Chor, Anna Mironov, Galina V. Beznoussenko, Alexander A. Mironov, Ella H. Sklan, George H. Patterson, Yoji Yonemura, Christoph Kaether, Koret Hirschberg

**Affiliations:** Department of Pathology, Sackler School of Medicine, Tel-Aviv University, Tel Aviv, Israel; Department of Physiology and Pharmacology, Sackler School of Medicine, Tel-Aviv University, Tel Aviv, Israel; Department of Clinical Immunology and Microbiology, Sackler School of Medicine, Tel-Aviv University, Tel Aviv, Israel; Institute of Protein Research, Russian Academy of Sciences, Moscow Region, Russian Federation; Bioinformatics Unit, G.S.W. Faculty of Life Sciences, Tel-Aviv University, Tel Aviv, Israel; IFOM, Fondazione Istituto FIRC (Fondazione Italiana per la Ricerca sul Cancro) di Oncologia Molecolare, Milan, Italy; Section on Biophotonics, NIBIB, National Institutes of Health Bethesda, Rockville, Maryland, USA; Leibniz Institute for Age Research - Fritz Lipmann Institute, Jena, Germany

## Abstract

COPII and COPI are considered to be analogous sets of vesicle coat protein heterocomplexes. Coupled to cargo selection, they mediate the formation of membrane vesicles translocating in opposite directions to differ rent destinations within the secretory pathway. Here, live cell and electron microscopy provided evidence for a different localization and mode of function of the COPII coat during protein export from the endoplasmic reticulum (ER). Pharmaceutical and genetic perturbations of ER-Golgi transport were used to demonstrate that COPII is recruited to membranes defining the boundary of ER-ER Exit Sites (ERES) where it facilitates selective cargo concentration. Uncoating of COPII membranes precedes cargo accumulation and fission of Golgi-bound carriers. Moreover, we report what may be direct transfer of cargo to the Golgi apparatus from Golgi-associated BFA sensitive ERESs. Finally, in *ldlF* cells the stably expressed functional ε-COPI-EYFP labeled both ERESs and anterograde carriers. These findings change our understanding of the role of coat proteins in ER to Golgi transport.

## Introduction

Secretory organelles exchange substantial fluxes of molecules via membrane carriers that translocate across the cytosol on microtubule (MT) tracks. To maintain organelle identity and functionality, the selection of the proteins and lipids that will travel in these carriers must be tightly regulated. Transport-competent proteins are those that are properly folded and contain transport signals. To be transported to its target organelle a protein has to be selected and concentrated within a well-defined membrane domain that will, in turn, mature into a transport vesicle or a carrier. The first sorting station for proteins in the secretory pathway are ER-exit sites (ERES), also called transitional ER (Balch et al., 1994). These are specialized membrane domains on the surface of the ER that are identified by the COPII heterocomplex and ER-Golgi recycling proteins such as ERGIC53 (Fujiwara et al., 1998). The COPII protein complex has been identified in yeast and in mammalian cells as a membrane coat that is involved in anterograde trafficking from the ER (Barlowe et al., 1994; Kirk and Ward, 2007). The complex formation is initiated by the Sar1 small GTPase which is recruited to the ER membrane by Sec12, a protein that acts as its GDP-to-GTP exchange factor (GEF). Sar1 recruitment triggers the assembly of Sec23/24 subcomplexes with cargo proteins, then Sec13/31 forms the outer cage of the coat. The assembling lattice deforms the membrane to become spherical or tubular membrane structures (Mancias and Goldberg, 2008; Zanetti et al., 2013). Mezzacasa et. al. demonstrated that the COPII-labeled transitional ER defines the quality control boundary after which misfolded proteins are not retrieved back to the ER (Mezzacasa and Helenius, 2002) (VSVGtsO45). In addition, they showed that the tsO45 thermoreversible folding mutant of vesicular stomatitis virus glycoprotein (VSVGtsO45) accumulated in COPII-labeled ERESs. We verified that cargo selection and accumulation was facilitated by COPII within dilated ERES in cells treated with BFA and Nocodazole (Dukhovny et al., 2008; Dukhovny et al., 2009).

Here, using living intact cells coexpressing fluorescent protein-tagged COPII and cargo proteins we followed ER to Golgi trafficking of VSVGtsO45. We found that carrier fission as well as the earlier step of cargo selection and accumulation of FP-VSVGtsO45 occurred essentially downstream from COPII-labeled membranes. Confocal and immunoelectron microscopy of cells treated with BFA and Nocodazole demonstrated that COPII defines the ER-ERES boundary where it facilitates cargo selection and concentration in ERES membranes. Using mutagenesis to perturb cargo binding of COPII we found that cargo accumulated in the ER thus supporting COPII localization and function at the ER-ERES boundary. Furthermore, we demonstrate that a subpopulation of COPI is localized to ERESs and is also seen on ER to Golgi anterograde carriers.

## Results

### The distribution of ER-exit sites

The distribution of ERES was analyzed in living intact cells coexpressing a COPII protein and ER markers tagged with fluorescent proteins (Fig1A). The COPII Sec24C-labeled ERES membranes were frequently associated with flat saccular-membrane elements of the ER as shown in figure 1A together with co-expressed Reticulon. In figure 1B, coexpression of mCherry-Sec24C with YFP-Rab1B depicts a subpopulation of ERESs that decorate the periphery of the Golgi apparatus.

**Figure 1.**
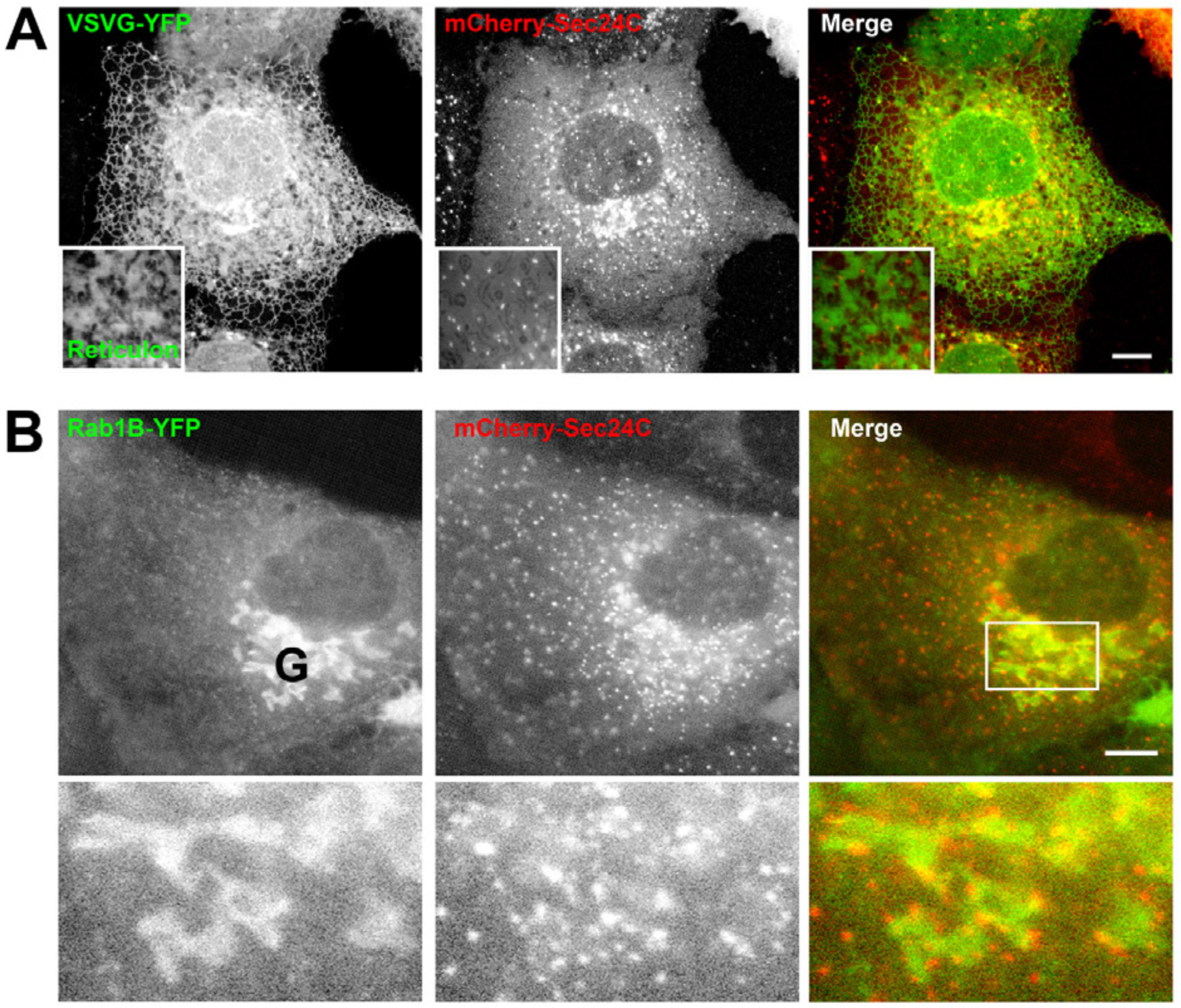
Intracellular distribution of COPII in living intact cells. **A.** Intracellular distribution of ERESs. A confocal image of a COS7 cell coexpressing the ER marker (at the non-permissive temperature 39.5°C) VSVG-YFP (green) or the ER membrane marker Reticulon-GFP (green in insert) and the COPII subunit Sec24C-mCherry (red). Bar = 5 µm. **B.** Colocalization of Sec24C-mCherry (red) and Rab1b-YFP. A confocal image of Huh7 cell coexpressing the Rab1b-YFP (green) and the COPII subunit Sec24C-mCherry (red). Golgi apparatus marked by G. Insert showing the Golgi apparatus is enlarged 3-fold below. Bar = 5 µm.

### Potential direct transfer of cargo from Golgi associated ERESs

Analysis of ER to Golgi transport of VSVG-YFP after shift to permissive temperature in Golgi associated ERESs demonstrated direct accumulation of the cargo in the ERESs-decorated Golgi membranes (Fig. 2A). To test the association of Golgi associated ERESs cells coexpressing Sec24C-mCherry and GalT-CFP were treated with BFA. Figure 2B demonstrates that the Golgi associated ERESs coincidently disappear with the Golgi membranes. This finding supports the prospect of direct association of ERESs with Golgi membranes. The possibility of alternative modes for secretory transport was recently raised suggesting temporal direct connections between organelles (Raote and Malhotra, 2019).

**Figure 2.**
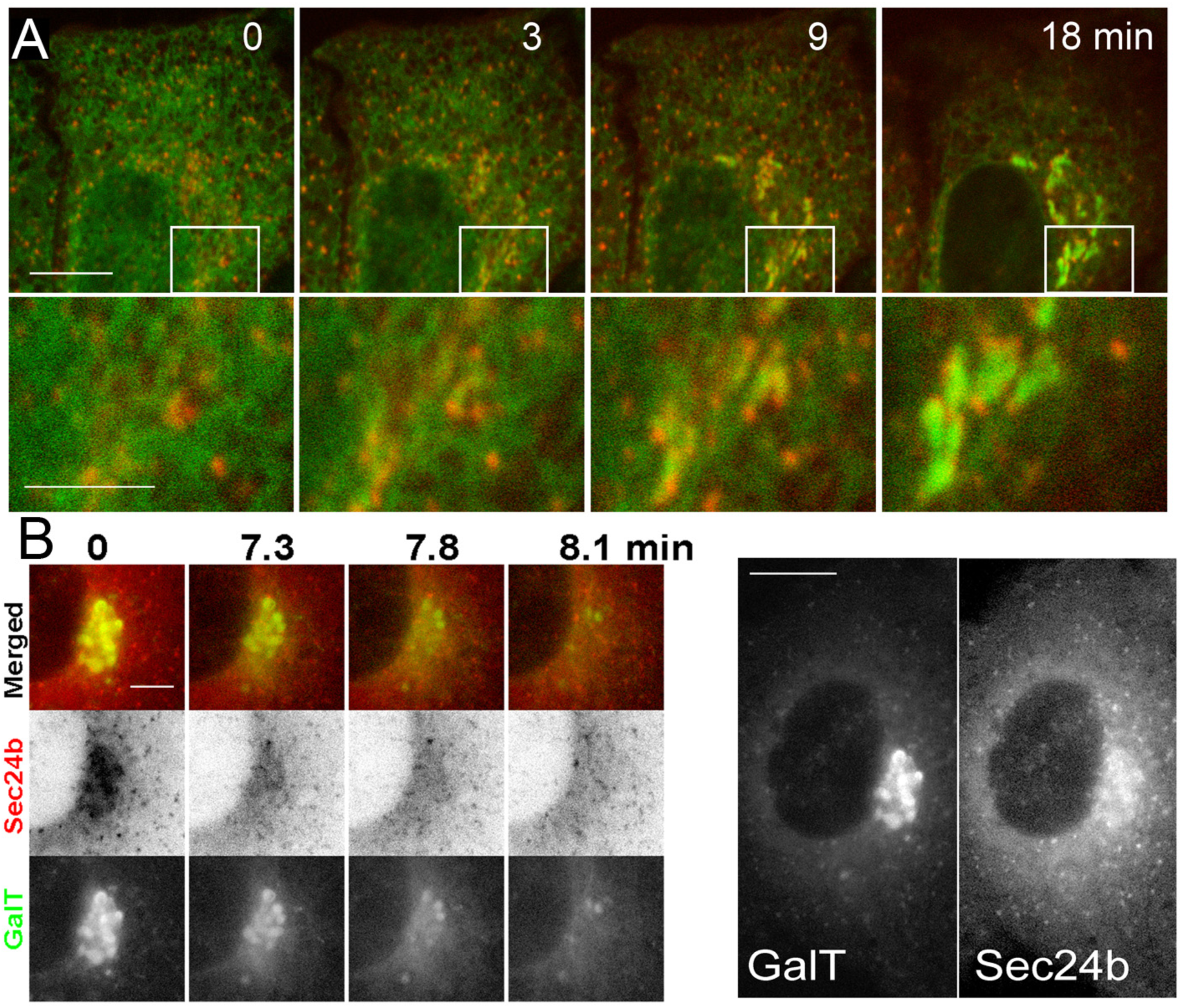
ER to Golgi transport in Golgi Associated ERES; Direct passage of VSVG-YFP (green) from Golgi-associated ERES to the Golgi apparatus. **A.** Time-lapse analysis showing an apparent direct passage of VSVG-YFP (green) from Golgi-associated ERESs to the Golgi apparatus in living cells coexpressing Sec24C-mCherry (red). Cells were treated and imaged as in b-c. Area in white rectangular border showing Golgi-associated ERESs is magnified 3.3-fold, Bar = 10 µm. **B**. Golgi-associated ERESs are disintegrated upon BFA-induced Golgi membranes blink-out. Representative images from a time-lapse sequence taken at 3 sec intervals of cells coexpressing a Golgi marker GalT-YFP (green and lower panel) and Sec24C-mCherry (red and inverted middle panel). Time points are after addition of 5µg/ml BFA. Entire cell is shown at the right-hand side, Bar = 10 µm.

### Dynamics of membrane-associated COPII

Next, we compared the distribution and dynamics of membrane-associated COPII with those of ER to Golgi vesicles and the growing plus ends of microtubules labeled with GFP-EB3. Figure 3A and the supplementary movie S3A show a maximum-intensity projection of 50 consecutive frames taken at 0.66 sec intervals. Compared to the rapid movement of transport carriers covering significant distances along microtubular tracks toward the Golgi apparatus, COPII-labeled membranes are essentially immobile. Similar results are obtained when COPII-labeled membranes are compared to microtubule plus-end tip progression using FP-EB3 (Fig. 3B and the supplementary movie S3B). While the growing microtubules covered distances of 20-25 micrometers in 3 minutes, COPII labeled membranes exhibited only limited random movements.

**Figure 3.**
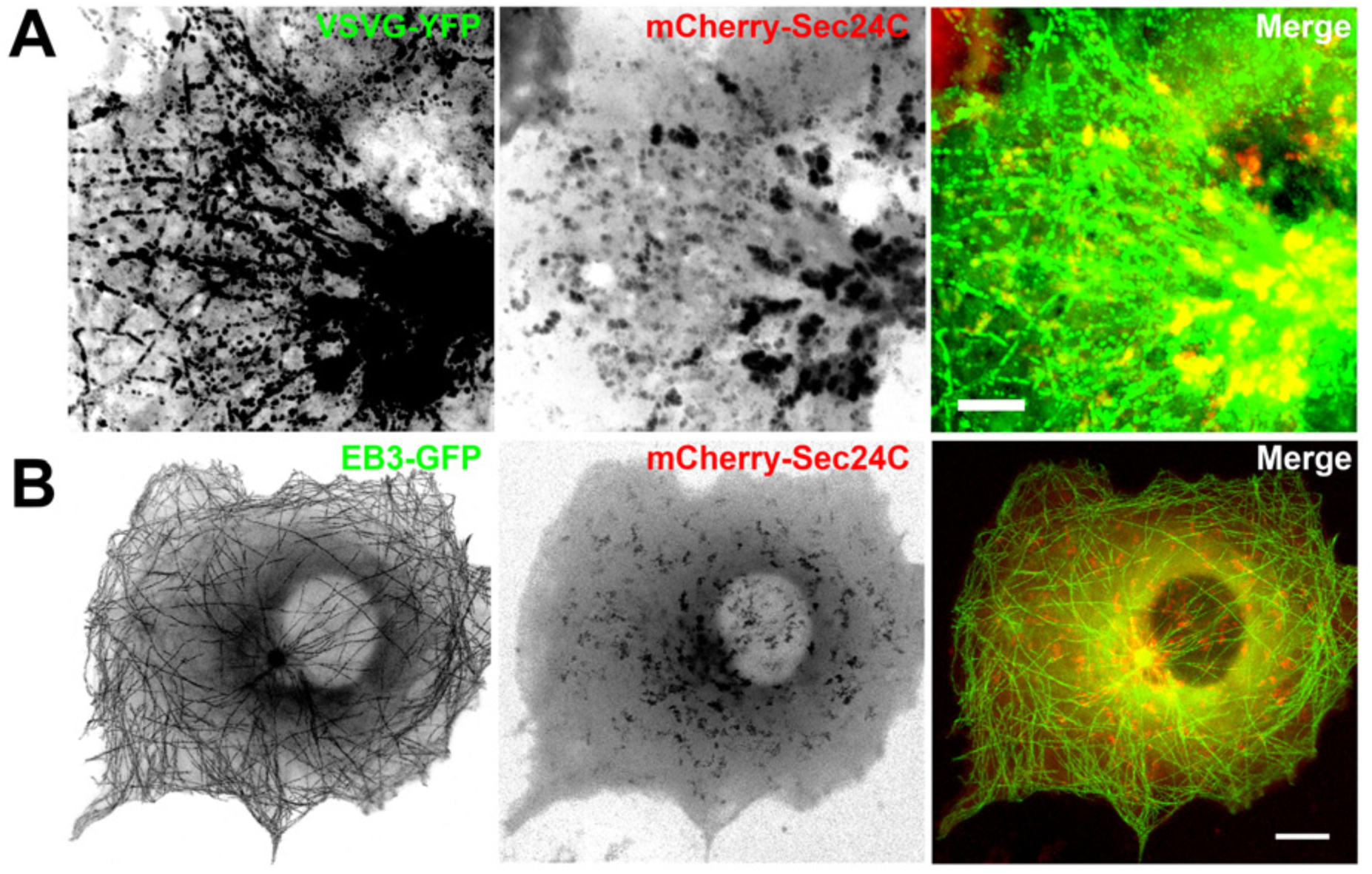
Dynamics of membrane-associated COPII. **A.** Comparison between membrane-associated COPII and ER to Golgi transport carriers. An inverted confocal projection of 50 images from a 0.66 sec interval time-lapse sequence of COS7 cell coexpressing VSVG-YFP (left and green) and the COPII subunit Sec24C-mCherry (center and red) after shift to permissive temperature 32°C is shown. Bar = 5 µm. See also supplementary movie S2A **B.** Comparison between membrane-associated COPII (middle and red) and the growing microtubule plus end marker EB3-GFP (left and green). An inverted confocal projection of 50 images from a 3.7 sec interval time-lapse sequence of COS7 coexpressing EB3-GFP (green) and the COPII subunit Sec24C-mCherry (red) is shown. Bar = 10 µm. See also supplementary movie S2B

### Cargo accumulation and subsequent fission occur downstream from immobile COPII coated membranes

Next, we analyzed the early stages of ER export of VSVG at a greater spatial and temporal resolution. ER export was analyzed at the level of a single ERES in living intact cells. To this end, cells coexpressing the cargo protein VSVG-YFP and Sec24C-mCherry were shifted to permissive temperature (32°C) after overnight incubation at the non-permissive temperature of 39.5°C. Figure 4A shows the dynamics of ER export in a living cell. Cargo is primarily concentrated and accumulated in growing dynamic tubular membranes that subsequently bud off. Based on these image sequences, ER export can be divided into two distinct consecutive stages. The first stage is the accumulation of cargo in ERES membranes (Fig. 4B-C). Subsequently, these membranes undergo fission to become membrane carriers (Fig. 4D-E). A key observation is that COPII uncoating occurred prior to both of the abovementioned processes, namely the early accumulation of cargo in ERES membranes and the subsequent fission. These data demonstrate a different mode of action to the current concept of cargo selection coupling to coated vesicle formation (Bethune and Wieland, 2018). A common explanation for the absence of membrane-bound COPII on forming carriers is that low levels of COPII are below the detection threshold. Thus, we tested whether we can detect as a positive control, another transport machinery protein that dynamically interacts with vesicle membranes. For this purpose, we used the small GTPase Rab1 tagged with mCherry. Cells coexpressing Rab1b-mCherry and VSVG-GFP were shifted to permissive temperature and ER to Golgi transport was followed using confocal microscopy. Figure 4F and supplementary movie S3D show the entire life history of an ER to Golgi carrier from its fission from an ERES through its translocation and to its fusion with Golgi membranes. The labeling of the carrier in the Rab1b-mCherry channel is unmistakably detectable throughout the entire pathway. These data argue against the view that membrane bound-COPII are below detection levels. Also, membrane bound Rab1 was suggested to replace COPII by the activity of the TRAPP-II complex to later recruit the ARF1 GEF GBF1 and COPI (Aridor, 2018). Finally, Fig. 4G demonstrate the release of GFP-tagged procollagen-I in cells coexpressing Sec24C-mCherry after temperature shift to 32°C and addition of ascorbic acid. Procollagen accumulates adjacent to the COPII labeled membranes and the two consecutive budding events occur on membranes without COPII. Thus, we hypothesize that COPII-coated membranes separate the ERES domain from the rest of the ER.

**Figure 4.**
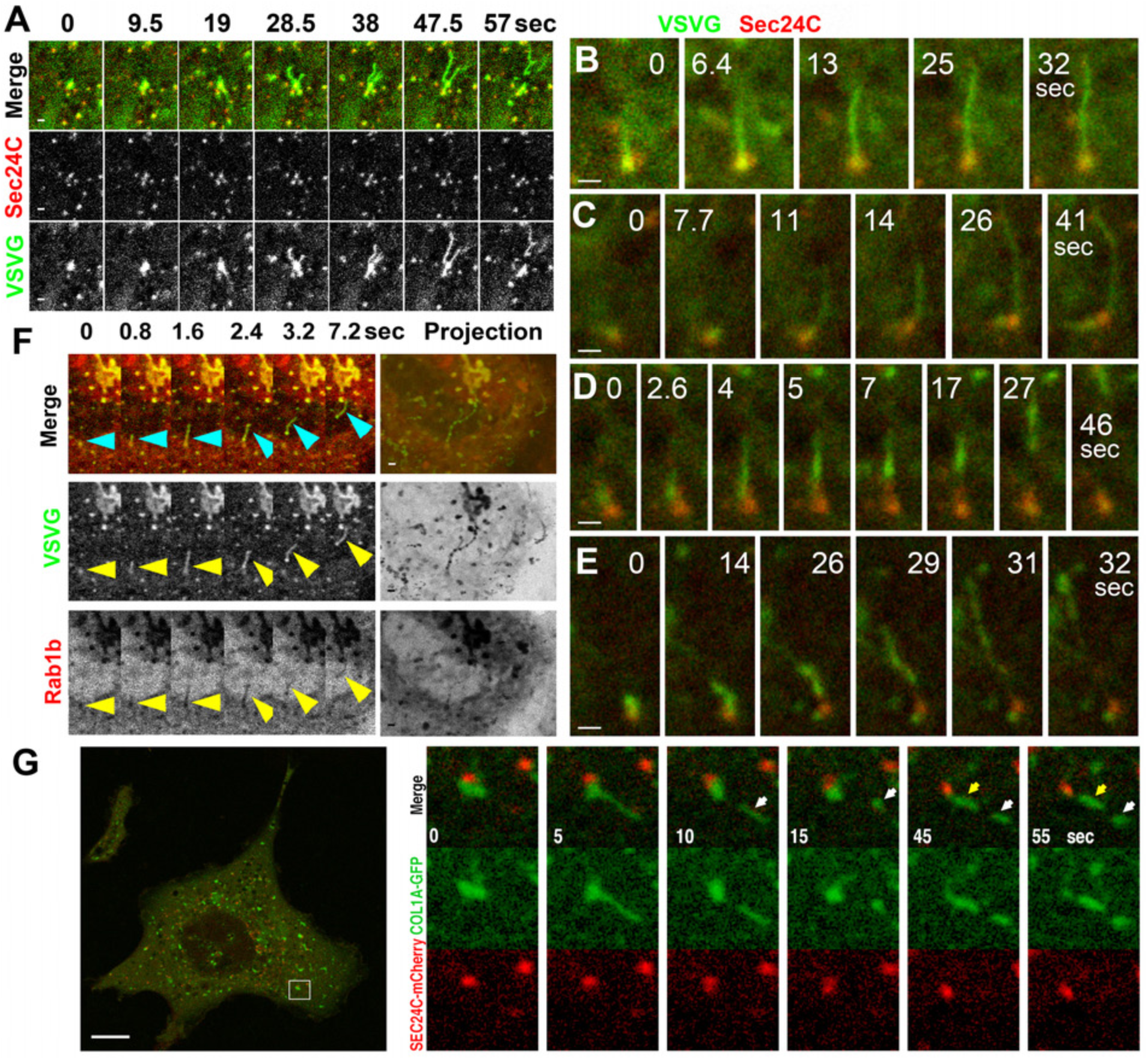
Cargo accumulation and subsequent fission occur downstream from immobile COPII coated membranes. **A.** Export of VSVG-YFP from the ER. Accumulation in ERES and fission of the cargo protein VSVG-YFP (green) in COS7 cells coexpressing Sec24C-mCherry (red). Bar = 1 μm. Cells were released from temperature-block during microscopy analysis at 32 °C after overnight post-transfection incubation at the non-permissive temperature 39.5 °C. Images were captured at 9.5 sec intervals. Bar =5 μm. See also supplementary movie S3A. **B-C.** Cargo accumulation in ERES membranes. COS7 cells were co-transfected with VSVG-YFP (green) and the COPII subunit Sec24C-mCherry (red). Images were captured at 2.6 sec intervals and treated as in A. Bar =1 μm. **D-E** Fission of transport carriers containing VSVG-YFP (green) from ERES in COS7 cells co-expressing Sec24C-mCherry (red). See also supplementary movies S3B through S3C. Bar =1 μm. **F**. Localization of the transport machinery protein Rab1-mCherry (red) in ER to Golgi carriers (arrowheads) containing VSVG-YFP (green) in Huh7 cells. Arrowheads point to carriers. Bar =5 μm. See also supplementary movie S3D. **G.** Two budding events in COS7 cells co expressing GFP tagged procollagen I (green) and Sec24C-mCherry (red). Cells incubated overnight in 39.5 °C were imaged after addition of ascorbic acid and shifting to 32 °C. white and yellow arrows show carriers after first and second budding, respectively. See also supplementary movie S3G. Bar =10 μm.

### COPII localizes to the ER-ERES boundary where it mediates selective cargo sorting and concentration in Nocodazole-BFA treated cells

Next, we established the relative localization of ERES-, COPII-coated- and ER membranes. It was previously reported that simultaneous inhibition of Arf1 activation by BFA (Lippincott-Schwartz et al., 1989) and microtubules polymerization by Nocodazole (Noc) (Hirschberg et al., 1998) preserves the cargo sorting functions of COPII, while blocking exit of cargo from the ER (Dukhovny et al., 2008). Release of VSVG-YFP at a permissive temperature in the presence of BFA and Noc results in active COPII-mediated sorting of VSVG-YFP into ERES membranes that now transform at the light microscope level into spherical structures (Fig. 5A) (Dukhovny et al., 2008). This perturbation of cargo export from the ER results in accumulation in ERES membranes that are adjacent to but spatially segregated from COPII-coated regions. Each cargo accumulation cycle culminates in rapid release of the cargo protein back into the ER. An identical pattern of localization of endogenous COPII under BFA/Noc treatment is shown by immunofluorescence analysis using anti-Sec24C antibody (Fig. 5B). To exclude unspecific side effects of the combined drug treatment the structures were demonstrated to be reproduced by replacing BFA/Noc by Golgicide A, a specific inhibitor of the Arf1-GDP exchange factor GBF1 (Saenz et al., 2009) and colchicine (Dubey et al., 2017), respectively (Fig. 5C). Three-dimensional reconstructions (Fig. 5D-F) including high resolution multiphoton structured illumination microscopy (Ingaramo et al., 2014) (Fig. 5E) demonstrate that COPII is bound to an elongated membrane under these conditions, similar to previously reported in *in-vitro* ultra-structural studies (Zanetti et al., 2013). Cells treated with BFA/Noc were analyzed using a variety of cargo proteins such as ERGIC53, GalT and cystic fibrosis transmembrane regulator (CFTR) (Fig. 5G-I), all of which showed identical pattern of accumulation in the apparently dilated ERES membranes and segregation from the COPII markers used. To confirm that the membrane and lumen of distended ERESs observed in BFA/Noc-treated cells are still continuous with those of the ER, we coexpressed VSVG-YFP and mCherry with a cleavable signal sequence conferring insertion into the ER lumen. Images of these markers (Fig. 5J and supplementary movie S5J) confirm that ERESs under BFA/Noc are linked to the ER. Furthermore, localization of membrane-bound COPII relative to both cargo-enriched membranes and the ER was determined using high speed confocal microscopy of BFA/Noc-treated cells. Fig. 5K and movies S5K1-2 contain deconvolved images demonstrating that COPII membranes are consistently positioned between the ER and ERESs. Arrows in the lower panel indicate the direction of flow of exported cargo. To investigate the ultrastructure of the apparently dilated ERESs we carried out electron-microscopy using immunogold anti-GFP antibodies on cells coexpressing Sec23C-GFP and VSVG-Scarlet. Fig. 5L shows representative images of the dilated ERESs. The immunogold labeling of Sec23 is concentrated at the area linking the structures to the ER. Moreover, the EM images demonstrate a highly fenestrated round hollow structure with a complex topology. The three-dimensional reconstruction of serial sections of one of these structures (Fig. 5M) confirms that the membranes are connected to form a structure previously reported for BFA treated cells as Glumerolini (Pavelka and Ellinger, 1993). Notably, as opposed to the reported structures the BFA/Noc structures revealed by EM had a significant inner cytosolic space probably resulting from its growth due to membrane and cargo inflow. Next, we analyzed the dynamics of COPII-mediated ER export functions in BFA/Noc-treated cells (Fig. 6). ERESs were monitored using a high-speed confocal microscope. As previously reported, ER exit sites undergo cycles of cargo accumulation culminating in rapid release of the cargo and membranes back into the ER (Fig. 6A, and supplementary movie S6A) (Dukhovny et al., 2008). In BFA/Noc-treated cells coexpressing the COPII subunit Sec24C and cargo, the levels of membrane-bound COPII fluorescence were apparently indifferent to the undulating concentrations of cargo within ERES (Fig. 6A). This result is consistent with COPII occupying the ER-ERES boundary. A complete single cargo accumulation cycle is shown in the form of a kymograph in Fig. 6B (see also supplementary movie S5B). Shown, is an increase in both ERES diameter as well as fluorescent intensity with time, indicating a net flow of both membrane and cargo from the ER through the COPII coated membranes. To demonstrate that COPII can distinguish between transport competent and incompetent proteins (Mezzacasa and Helenius, 2002) in BFA/Noc-treated cells, we compared COPII-mediated sorting of wildtype and the Δ508 mutant of CFTR (Fig. 6C) (Wang et al., 2004). While wildtype CFTR accumulates in ERESs, the misfolded Δ508 mutant is excluded.

**Figure 5.**
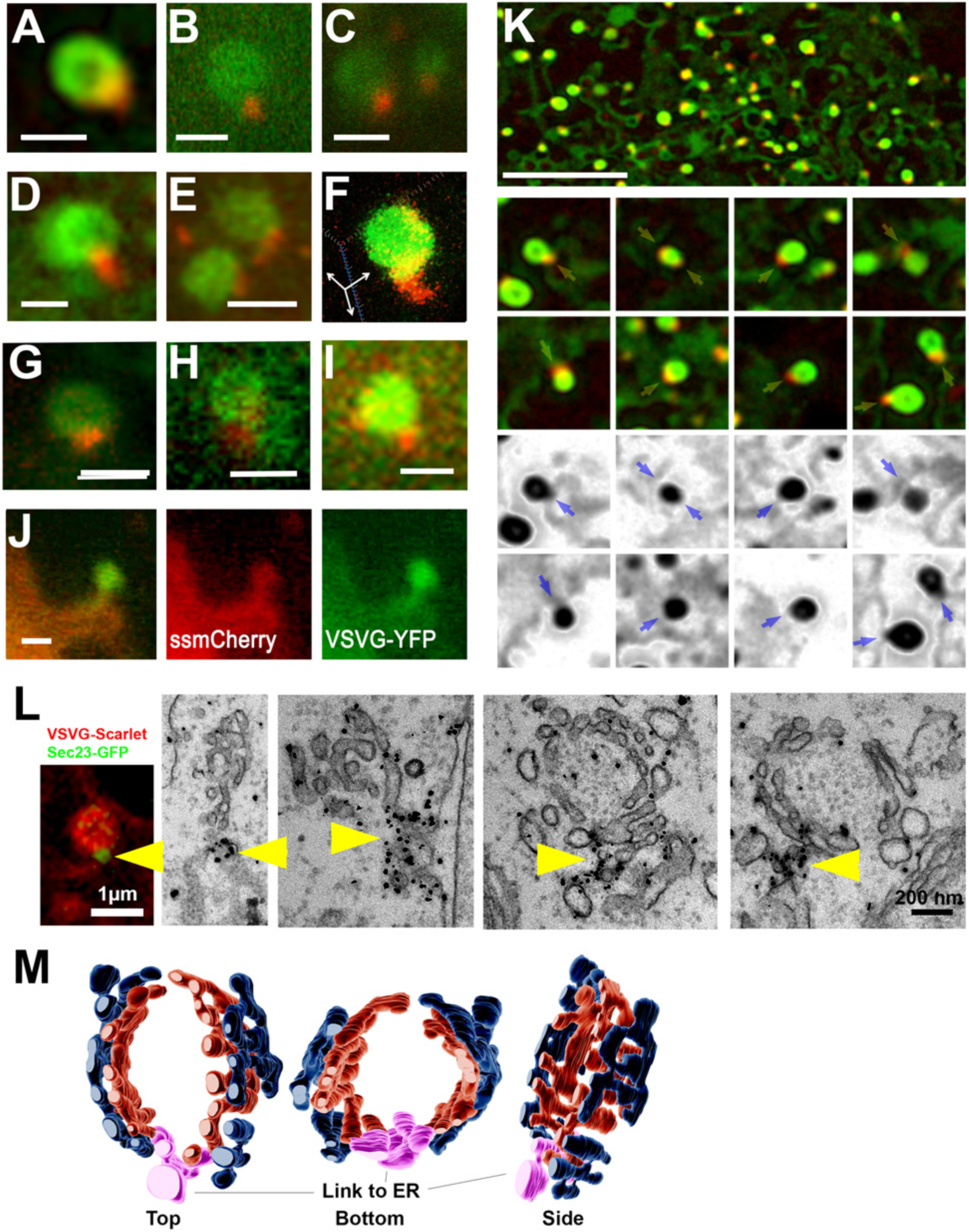
Localization of COPII to the ER-ERES boundary in BFA/Noc-treated cells. **A-F** Relative localizations of Sec24C-mCherry (red) and VSVG-YFP (green) within representative ERESs of COS7 cells under BFA/Noc treatment. Bars = 1 µm **A.** Confocal image of a single ERES. **B.** Immunofluorescence analysis of endogenous Sec24C (red) in a cell expressing VSVG-YFP (green) under BFA/Noc treatment. **C.** An ERES of a cell under Golgicide A and colchicine treatment. **D.** A three-dimensional deconvolved image of a single ERES. **E.** A high resolution three-dimensional reconstruction obtained using MSIM. **F.** A three-dimensional reconstruction of confocal thin z-section series **G.** Relative localizations of Sec24C-mCherry (red) and ERGIC53-YFP (green) within a representative ERES of COS7 cells under BFA/Noc treatment. Bars = 1µm. **H**. Relative localizations of Sec24C-mCherry (red) and GalT-YFP (green) within a representative ERES of COS7 cells under BFA/Noc treatment **I**. Relative localizations of Sec24C-mCherry (red) and CFTR-YFP (green) within a representative ERES of COS7 cells under BFA/Noc treatment **J.** ERES is connected to the ER via a tubular-shaped membrane in a living cell coexpressing VSVG-YFP (green) and the ER luminal marker signal sequence-mCherry (red) under BFA/Noc-treatment. Bar = 1 µm (See also supplementary movie S4J) **K.** The COPII-bound membranes are localized between ERES and ER. Deconvolved confocal image series from living intact COS7 cells coexpressing Sec24C-mCherry (red) and VSVG-YFP (green) in the presence of BFA/Noc. Yellow and blue arrows in the inverted images point to ER-ERES COPII–coated intersections. The VSVG channel is shown inverted below. (See also supplementary movies S4K1-2). **L.** Immunogold electron microscopy with anti-GFP antibodies of ERESs in BFA/Noc-treated cells. COS7 cells expressing Sec23-GFP (green) and VSVG-Scarlet (red) under BFA/Noc treatment were fixed after 30 min at permissive temperature 32°C. Yellow arrowheads point to COPII at sites of contact to ER. Bar = 200nm. **M.** Tomogram of a serial section from L. showing connectivity of all the membranes upstream to the COPII.

**Figure 6.**
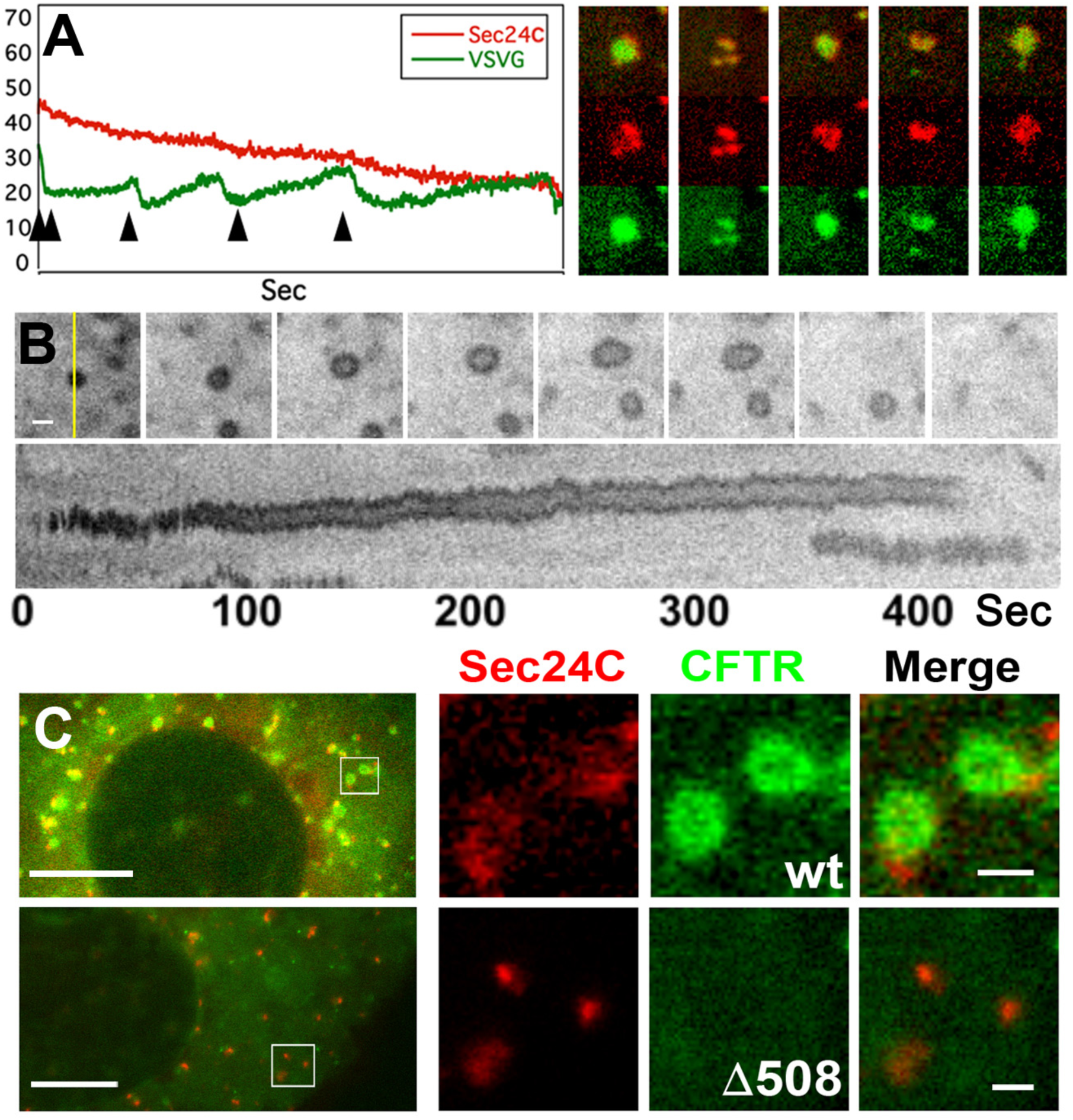
COPII-mediated cargo sorting dynamics in BFA/Noc-treated cells. Analysis of COPII-mediated cargo concentration dynamics in ERESs of BFA/Noc-treated COS7 cells. Cells coexpressing VSVG-YFP (green) and Sec24C-mCherry (red) were imaged at 0.2 sec intervals. The fluorescence intensities of VSVG-YFP (green line) and Sec24C-mCherry (red line) in an ERES are plotted and representative images are shown on the right end side. Time-points of images is shown by black arrowheads in the graph. **B.** Selected images and kymograph demonstrating cargo and membrane flow into ERESs in BFA/Noc-treated COS7 cells expressing VSVG-YFP. Kymograph is generated from the thin section marked with the yellow line. Bar = 1 µm. **C.** The misfolded mutant cargo protein CFTRΔ508-GFP (green, bottom) is excluded from ERES in BFA/Noc-treated COS7 cells. Cells are coexpressing Sec24C-mCherry (red) with either wild type CFTR-GFP (green, top) or CFTRΔ508-GFP (green, bottom). Insets are enlarged 5 fold. Bar on the left panels = 10 µm. Bar on the right panels = 1 µm.

### Mutagenesis of the cargo-binding-Sec24B subunit disrupts cargo accumulation in ERESs causing ER retention

Sec24 is a COPII subunit that directly interacts with cargo (Wendeler et al., 2007). The B isoform of Sec24 interacts with acidic export motifs found in numerous surface proteins including VSVG and CFTR (Wang et al., 2004). To enhance the binding between Sec24B and the di-acidic motif of VSVG, we used structure-based computer modeling of Sec24B (PDB ID 3EH1) (Mancias and Goldberg, 2008) and designed a mutant of Sec24B by substituting Valine at position 932 with the basic Arginine (V932R). This results in an additional positive charge in the binding pocket for acidic export motifs of cargo proteins (Fig. 7A-B). A stronger binding of the VSVG tail by the V932R mutant compared to the wildtype was predicted by both allowed docking peptide states and surface electrostatic potential analysis (Dolinsky et al., 2007; London et al., 2011; Pettersen et al., 2004). Primarily, we confirmed that the mutant Sec24B_V932R_ bound VSVG using co-immunoprecipitation (Supplementary Fig. S1). Fluorescence recovery after photobleaching was measured for wildtype or mutant Sec24B in BFA/Noc-treated cells coexpressing VSVG (Fig. 7C). As predicted by the abovementioned models in Fig. 7A-B, the results show that membrane-bound Sec24B_V932R_ has a 1.5-fold slower on/off rate and a smaller mobile fraction compared to wildtype Sec24B, consistent with a stronger binding of cargo by Sec24B_V932R._ Expression of Sec24B_V932R_ blocked VSVG transport causing its accumulation in the ER after release in permissive temperature (Fig. 7D-F). This accumulation in the ER supports our previous results on the localization of membrane-bound COPII to the ER-ERES boundary, where it controls access of ER proteins to ERES. The phenotype of Sec24B_V932R_ was further studied using CFTR, a di-acidic-motif-containing polytopic surface protein(Wang et al., 2004). Fig. 6H shows that in BFA/Noc-treated cells GFP-CFTR is excluded from ERESs in cells expressing Sec24B_V932R_ (Fig. 7G). The data here demonstrate that COPII localize and functions at the ER-ERES boundary, essentially mediating selective cargo concentration from ER through COPII coated membranes to ERES.

**Figure 7.**
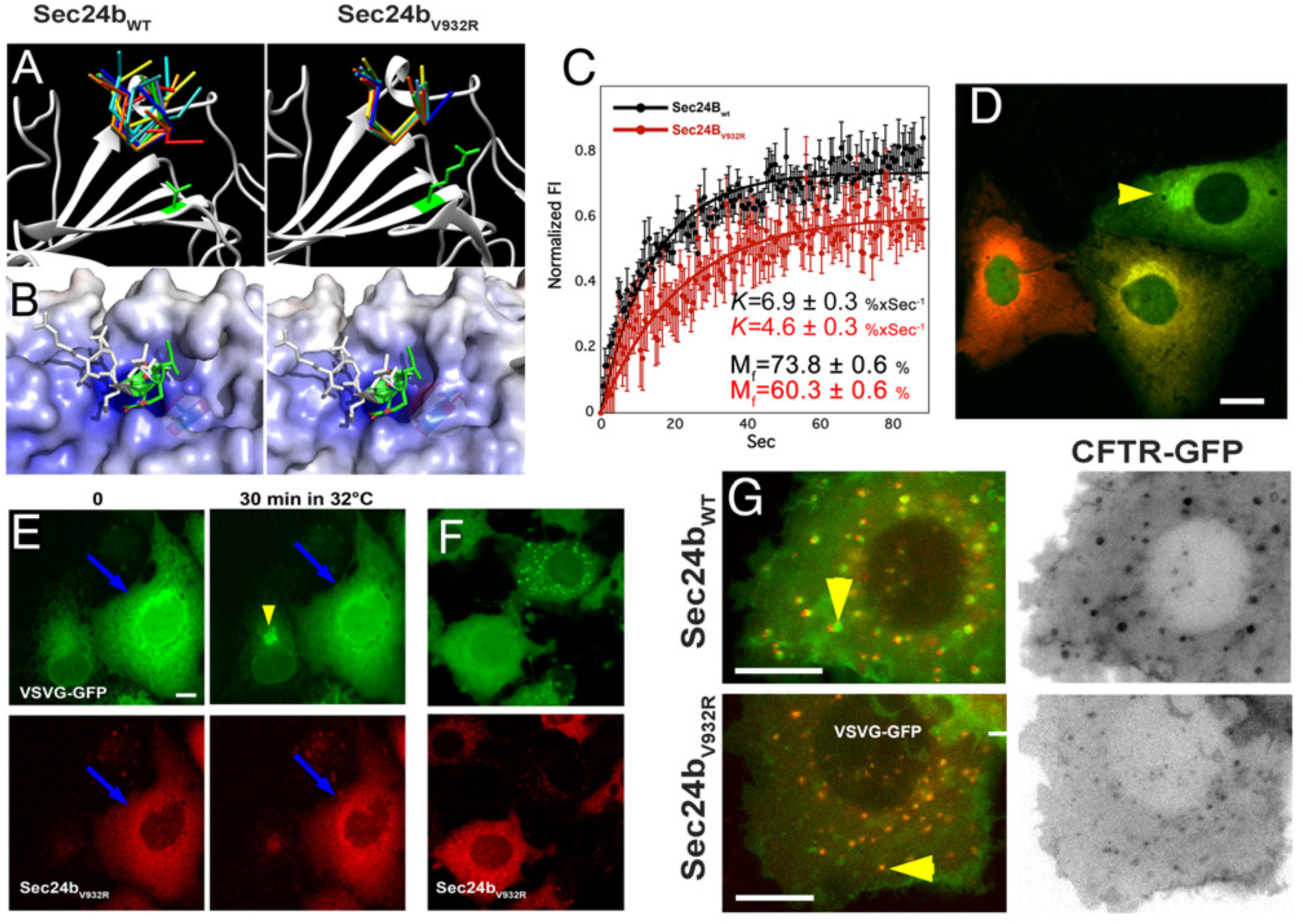
Perturbation of the COPII-cargo interface using mutagenesis of Sec24B. **A-** Computer modeling based on x-ray structure of Sec24-VSVG tail complex is shown for wildtype and for Val to Arg replacement at position 932. **A**. Allowed conformations of VSVG tail. Replaced amino acids with their structural representations are shown in green **B**. Electrostatic potential and solvent accessibility maps show a more definite pocket for the VSVG tail binding site in the V932R mutant. **C**. Fluorescence recovery after photobleaching of wildtype (black) and mutant mCerry-Sec24B (red) coexpressed with VSVG-YFP in BFA/Noc treated cells as described in the Methods section. The mutant Sec24B_V932R_ has a smaller mobile fraction as well as a slower recovery rate suggesting a stronger binding of the VSVG cargo. **D**. Dose-response of overexpression of the Sec24B_V932R_ (red) mutant inhibiting ER export of VSVG-YFP (green). Image shows three cells with different levels of expression of Sec24B_V932R_ with a corresponding degree of ER export inhibition (time point after 32°C). The Arrowhead points to the Golgi apparatus. **E**. Shifting VSVG-YFP (green) to permissive temperature in cells coexpressing the mutant Sec24B_V932R_ (red) results in ER retention. The yellow arrowhead points to the Golgi apparatus in a cell not expressing Sec24B_V932R_. Blue arrow points to cells with VSVG retained in the ER **F.** Expression of the Sec24B_V932R_ mutant (red) in BFA/Noc-treated cells results in ER retention of VSVG-YFP (green). Yellow arrow point to a cell that does not express the Sec24B_V932R_ showing normal accumulation in ERES. **G.** The Sec24B_V932R_ mutant (red) but not the wild type blocks sorting of CFTR-GFP (green and inverted panel on right) in BFA/Noc cells. Yellow arrowheads point to ERESs.

### Evidence for the role of COPI in anterograde trafficking

The localization of COPII to the Boundary between the ER and ERES membranes prompts a question as to the identity of the coat protein for anterograde ER to Golgi carriers. Although it is accepted that COPI is associated with retrograde transport there are several observations that clearly show the opposite: The localization of COPI on anterograde vesicles or carriers has been previously demonstrated (Presley et al., 2002). A sequential mode of action was demonstrated by scales et. al. and Shima et al. where COPI was shown to replace COPII on ER to Golgi vesicles (Scales et al., 1997; Shima et al., 1999). Similar results were reported by Stephens et al. (Stephens et al., 2000) using microinjected anti-COPI antibodies to label ER to Golgi vesicles. In our hands using confocal time lapse microscopy of living intact LdlF cells (Presley et al., 2002) coexpressing Sec24C-mCherry and the ER marker KDEL-BFP2 we found that COPI localized to the cytosol as well as to Golgi apparatus membranes and to ERESs. Also, ε-COP-YFP appears on anterograde Golgi-bound membrane carriers (Figure 8)

**Figure 8.**
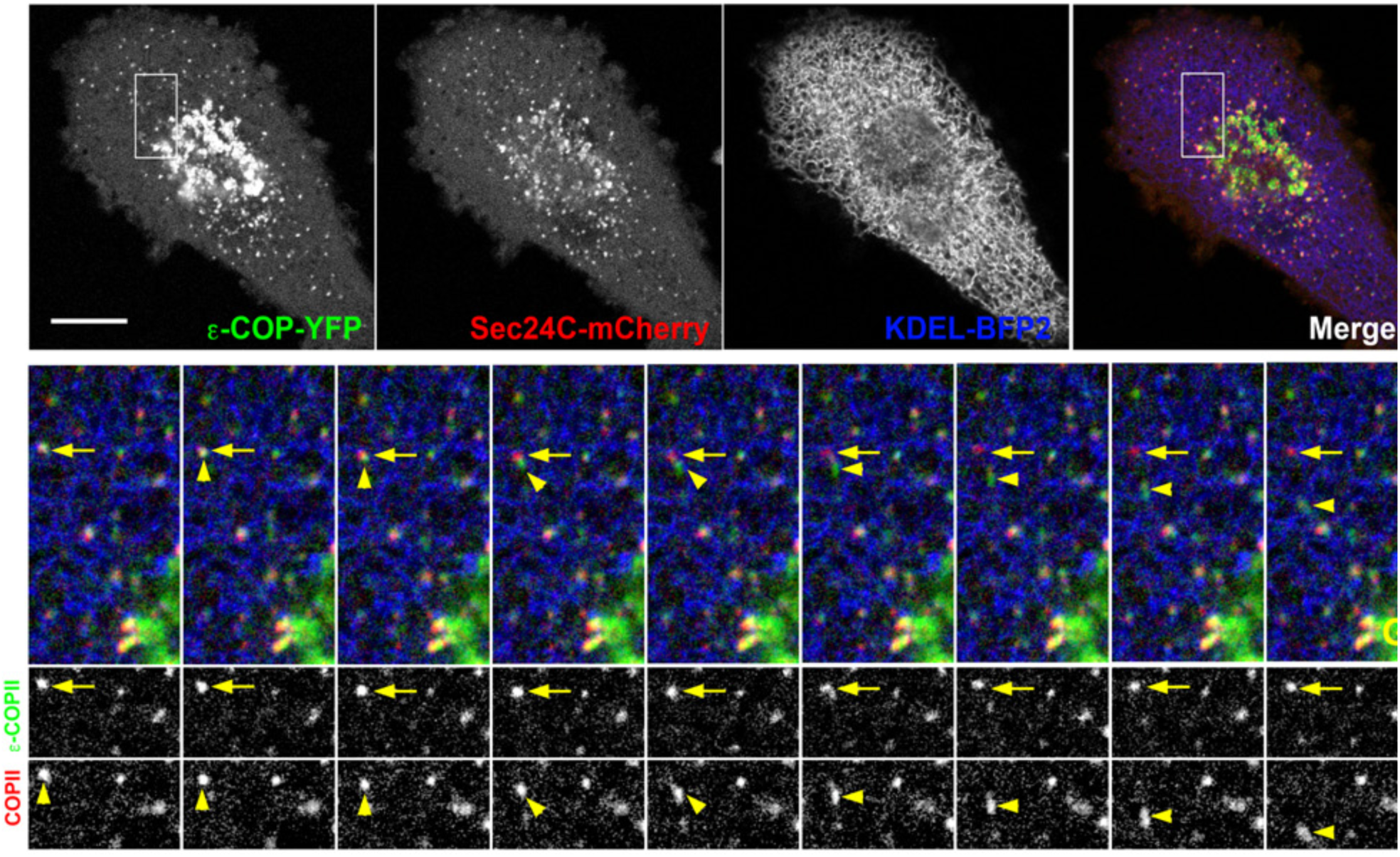
Colocalization to and translocation from ERESs of COPI coated membranes. Time lapse microscopy analysis of living LDLF cells stably expressing ε-COP-YFP (green) cotransfected with Sec24C-mCherry (red) and KDEL-BFP (Blue). Yellow arrows point to ERES labeled with Sec24C and yellow arrowheads point to COPI-membranes moving towards the Golgi (G) Bar = 10 μm.

## Discussion

Our findings are summarized in the scheme in Fig. 9: COPII mediates the sorting of transport-competent proteins by dynamic binding to stable domains of elongated membranes that comprise the ER-ERES boundary. Previous studies demonstrated that COPII can form a boundary for the ER quality control machinery (Mezzacasa and Helenius, 2002). A recent finding, consistent with these data, established that COPII is fundamental for ER retention of transport-incompetent proteins (Ma et al., 2017). The data presented in this paper provides a novel mode of action for the COPII complex. The conventional vesicle coat model of COPII is based on ultrastructural studies of purified COPII components and artificial membranes (Bethune and Wieland, 2018; Mancias and Goldberg, 2008). These studies could not identify the function of COPII at the ER-ERES boundary as vesicle formation was analyzed in a cell free system that cannot assign a precise intracellular localization to the process. The use of semi-permeable cells to reconstitute COPII vesicle formation may also have altered the fragile tubular-vesicular structure of the ERESs due to potentially destructive effects of the permeabilizing detergent on membranes. (Supplementary figure S2). To this end, the concern that a substantial amount of data on COPII vesicles is indirect, has been raised previously (Mironov, 2014; Mironov et al., 2003). Recent ultrastructural studies, demonstrated and analyzed (O’Donnell et al., 2011; Zanetti et al., 2013) COPII-mediated tubular membrane structures. These *in-vitro* structures may correlate with the *in-situ* localization of COPII to a membrane bottleneck of the ER-ERES boundary. Our study is based in part on cells treated with BFA and nocodazole. We argue that despite the morphological changes and block of vesicle formation, COPII does not change its localization at the ER-ERES boundary. This was further confirmed by both light and electron microscopy analysis. Moreover, the electron microscopy images revealed a topologically complex structure of ERES under these conditions. Essentially, under BFA/Noc treatment ERES membranes are spherical in shape but are highly fenestrated yielding a spherical superficial network of tubular membrane with a hollow cytosolic center. It remains to be studied how these complex membranes are formed as well as the identity of proteins involved. The data presented here lays the foundation for an alternative mechanism of how the COPII complex orchestrates the functions of the ERES domain. Essentially, COPII mediates the selection and concentration of cargo within ERESs. We propose two distinct mechanisms that together facilitate the selective cargo entry into ERESs. The first is based on interaction of cargo protein with its surrounding membrane. It is widely accepted that an ER-Golgi-PM gradient of membrane thickness is key for targeting protein in the secretory pathway (Sharpe et al., 2010). Thus, In the ER, the transmembrane domain of PM-targeted integral proteins is longer than the average thickness of the ER membrane. Alleviation of this mismatching can thermodynamically drive cargo membrane proteins movement from the ER into ERESs. Indeed, shortening the TMD length of VSVG slows its accumulation and decreases its concentration in ERESs as well as in the trans Golgi network thereby affecting the dynamics of its passage through the secretory pathway (Dejgaard et al., 2008; Dukhovny et al., 2009). However, the cargo TMD-lipids hydrophobic mismatch predominantly functions in ERES retention since the membrane tension caused by the mismatching is only local. It is yet to be directly demonstrated that the membrane deformation resulting from such hydrophobic mismatch provides the necessary driving force for this process. The second mechanism for cargo entry into ERESs is based on the anticipated net directional ER to ERES movement of the membrane-bound COPII coat. This ER to ERES directed flow is generated by exclusive recruitment of COPII at the ER side. This occurs simply due to the restricted localization of the COPII recruiting small GTPase Sar1 to the ER membrane side (Kurokawa et al., 2016). Thus, the Sec24 subunit of the treadmilling COPII, interacts and drags the cargo from the ER towards the ERES (Nishimura and Balch, 1997; Wang et al., 2004). Together, these two mechanisms result in the capacity of the COPII complex to generate a selective and a directed flux of cargo and membrane to the ERES as shown in Fig. 7. When considering the well-studied vesicle formation-coupled cargo selection by COPII, it can be envisioned that this novel alternative model is a variation on the theme resulting from an early pre-budding uncoating of COPII from the membranes allowing more on-demand flexible and extensive proliferation of membrane to later generate large ER to Golgi carriers (Presley et al., 1997). Also, the recent idea of transport in non-vesicular tubular elements (tunnels) suggested by Raote and Malhotra is consistant with the localization of COPII reported here and is consistant with the direct transport from Golgi associated ERESs in Fig. 2 (Raote and Malhotra, 2019). In yeast and plants, ERESs facilitate direct transport of cargo to the Golgi apparatus (Kurokawa et al., 2014; Langhans et al., 2012). The early uncoating of COPII may also resolve the issue of how large cargo molecules or complexes can be accommodated in ER-derived vesicles (Mironov et al., 2003). This model is also consistent with recent work demonstrating the ER export of procollagen (Raote et al., 2018). It is less clear what is the repertoire of proteins that binds ER exit site membranes after uncoating of COPII. A key protein in this context is Rab1 which is activated and recruited by the TRAPP GEF complex and recruits GBF1 to ERES membranes which successively recruits the COPI complex via ARF1 activation. The exact timings of these processes is not clear however we can detect Rab1 from immediately after COPII uncoating until arrival and fusion of the carriers with the Golgi apparatus. The early recruitment of COPI to ERESs and ER to Golgi carriers reported previously and here is consistent with our data and provides a straightforward explanation why BFA blocks ER export. In sum, we present evidence for an extended model for the mechanism of the COPII sorting function. This model is of major significance as it allows a better understanding of mechanisms of protein retention and export from the ER in heath and disease.

**Figure 9.**
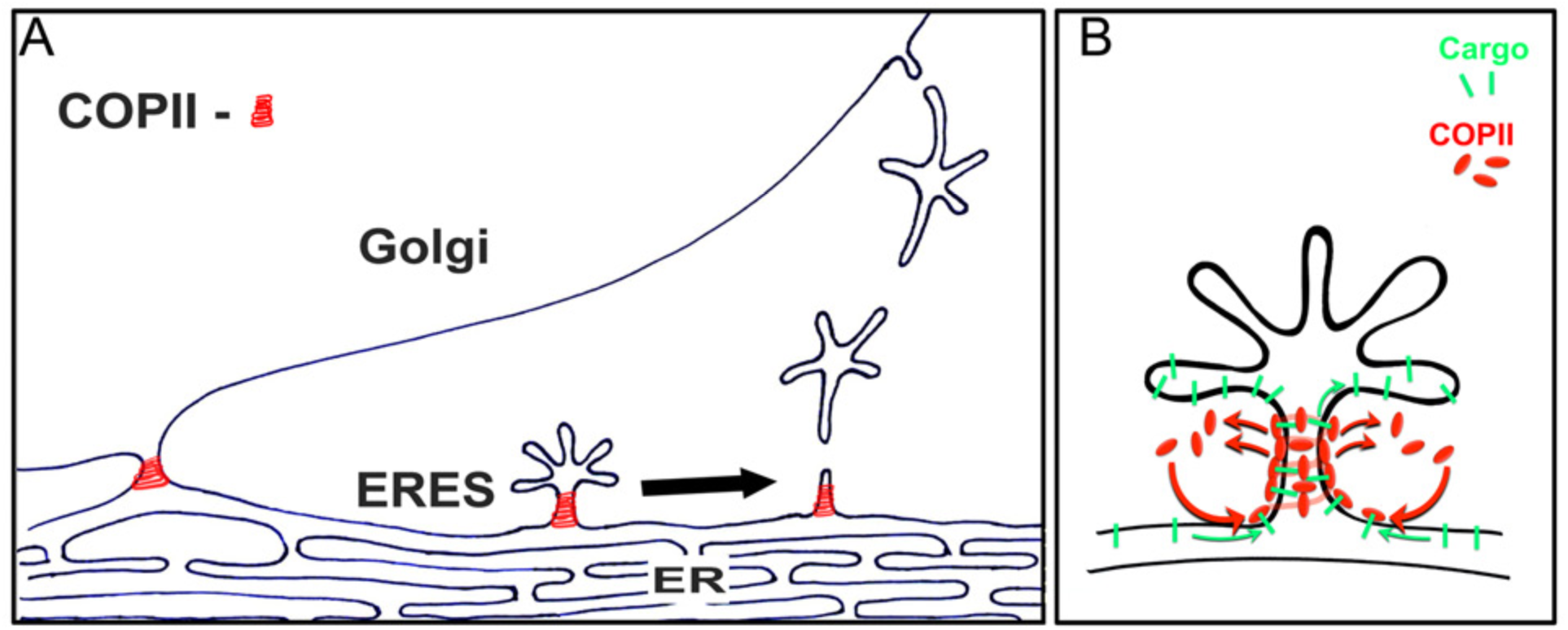
Schematic representation for the localization and function of the COPII heterocomplex at the ER-ERES boundary. **A.** COPII dynamically binds stable domains of elongated tubular membranes that comprise the ER-ERES boundary. Both cargo accumulation and fission occur on COPII-free membranes downstream to the coated membranes. Golgi associated ERESs may facilitate direct transfer of cargo from ER to Golgi membranes. **B.** A detailed model of the dynamic COPII cargo sorting machinery. Recruitment and assembly of the COPII coat is initiated exclusively at the ER side, generating a directed treadmilling movement of COPII, its bound cargo and surrounding membrane towards the ERES membrane.

## Methods

### Reagents and constructs

All reagents were purchased from Sigma Chemical Co. (St Louis, MO) unless otherwise stated. Human Sec24B was subcloned into pmCherry-C1 (Clontech) using *Sal*I and *Bgl*II restriction sites and verified by sequencing. Human Sec24C was subcloned into pmCherry-C1 or pEYFP-C1 (Clontech) using *Xho*I and *Sac*II restriction sites and verified by sequencing. Human Rab1B was cloned into pEGFP-C1 (Clontech) using *Xho*I and *BamH*I restriction sites and verified by sequencing. ss-mCherry (hen egg lysosime signal sequence. YFP-VSVGtsO45 was prepared as described elsewhere (Ward et al., 2001). GalT-YFP was prepared as described elsewhere(Zaal et al., 1999). GFP-CFTR and CFTRΔ508-GFP were a kind gift from R. Kopito (Stanford, CA). Rab1b was prepared as described elsewhere (Nevo-Yassaf et al., 2012). Reticulon-GFP was a kind gift from T. Rapoport (Boston, MA), CPE ws a kind gift from R. Harbefeld (Tel-Aviv, IL). Sec24B V932R mutant was prepared using the Quick-Change kit from Stratagene (La Jolla, CA). The primer used for the PCR reaction was: 5’-CAAGAAAAATTGGGTTTGAAGCTAGAATGAGAATAAGGTGTACTAAAGG-3’.

### Cell culture and transfections

COS-7, Huh7 or HeLa cells were grown at 37°C in a 5% CO_2_–humidified atmosphere. Cell cultures were maintained in Dulbecco’s modified Eagle’s medium (DMEM) supplemented with 10% (v/v) fetal bovine serum and penicillin and streptomycin (Biological Industries, Bet-Haemek, Israel). A final concentration of 1% (vol/vol) nonessential amino acids was added to the Huh7 cells culture media. Polyethyleneimine ‘MAX’ transfection reagent (Warrington, PA, USA), was used following the manufacturers’ protocols for plasmid DNA transfections of subconfluent COS7 and Huh7 cells. Confocal laser scanning microscopy experiments were carried out from 18 to 24 h after transfection.

### Live-cell microscopy

Cells were imaged in DMEM without phenol red but with supplements, including 20 mM HEPES, pH 7.4. Transfection and imaging were carried out in a 35-mm glass-bottomed microwell dish (MatTek, Ashland, MA) or on glass coverslips. A Zeiss LSM700 or LSM800 confocal laser-scanning microscope (Carl Zeiss MicroImaging, Jena, Germany). Fluorescence emissions resulting from 405-nm excitation for CFP 488-nm excitation for GFP and 543-nm excitation for mCherry were detected using filter sets supplied by the manufacturer. The confocal and time-lapse images were captured using a Plan-Apochromat ×63 1.4-numerical-aperture (NA) objective (Carl Zeiss MicroImaging). The temperature on the microscope stage was held stable during time-lapse sessions using an electronic temperature-controlled airstream incubator. Images and movies were generated and analyzed using the Zeiss LSM Zen software and NIH Image and ImageJ software (W. Rasband, National Institutes of Health, Bethesda, MD). High frequency images were captured using a Nikon Ti microscope equipped with Yokogawa CSU X-1 spinning disc, controlled by Andor IQ2 software or a spinning disk confocal unit (CSU-X1; Yokogawa electric corporation) attached to an Axio Observer Z1 microscope (Carl Zeiss).

### Confocal lasers scanning microscopy, time-lapse imaging and FRAP analyses

For FRAP measurements, a ×63 1.4-NA Plan-Apochromat objective was used on an inverted LSM800 system. Photobleaching of GFP was performed using four to six rapid scans with the laser at full power. Pre- and postbleach images were captured at 0.5- to 3-s intervals, using low laser intensity. Fluorescence recovery in the bleached region during the time series was quantified using LSM Zen software (Carl Zeiss MicroImaging). For presentation purposes, 16 bit confocal images were exported in TIFF, and their contrast and brightness were optimized in Adobe Photoshop software (San Jose, CA) or in ImageJ (Wayne Rasband, NIH, Bethesda MD). The characteristic fluorescence recovery time (τ) values for membrane-bound Sec24b turnover were calculated from the photobleaching data by fitting the data to a simple exponential equation using Kalaidagraph software (Synergy Software, Essex Junction, VT).

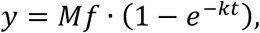

*Mf*= mobile fraction. R^2^ = 0.98 and 0.97 for wt and mutant, respectively.

### Immunofluorescence antibody staining

Cells cultured on coverslips were fixed at room temperature in a mixture of 2% paraformaldehyde–PBS, for 10 min, then washed three times with 1× PBS + 3% FCS. Cells were permeabilized and labeled simultaneously by incubation with appropriate primary antibodies: anti-Sec24C (4ug/ml, Abcam catalog no. Ab122635) and 0.01% saponin at room temperature for 1 h. cells were washed for three times with 1× PBS for 5 min at room temperature and incubated for 1 h at room temperate with appropriate Cy3 secondary antibodies (dilution 1:200, Jackson ImmunoResearch, West Grove, PA) and 0.1% saponin. Images were acquired using a Zeiss Pascal confocal laser-scanning microscope as described above.

### Electron microscopy

COS7 cells expressing Sec23-GFP and VSVG-Scarlet under BFA/Noc treatment were fixed 30 min after shift to permissive temperature for immuno-EM(Beznoussenko et al., 2014) Cells were then incubated with polyclonal anti-GFP antibody (Abcam, UK) for 2 hours and subsequently with anti-rabbit Fab’ fragment nanogold conjugates (NanoProbes) for 2 hours, enhanced then with GoldEnhance kit (NanoProbes) according to the manufacturer instruction. Epon embedding and sectioning of the gold-labeled cells, two-step electron microscopic tomography and three-dimensional reconstruction were performed as reported earlier(Beznoussenko et al., 2014; Beznoussenko et al., 2016). Sections were analyzed under Tecnai-12 electron microscope (ThermoFisher, Einhoven, The Netherlands) equipped with Analysis software.

### Immunoprecipitation and western blot analysis

Twenty-four hours following transfection, cells were washed with phosphate-buffered saline (PBS) and solubilized in lysis buffer (50 mm Tris-HCl, pH 7.4, 1% NP-40, 0.25%sodium-deoxycholate, 150 mm NaCl, 1 mm EDTA) containing protease inhibitor cocktail (Sigma). Extracts were clarified by centrifugation at 12 000 *g* for 15 min at 4 °C. Following SDS– polyacrylamide gel electrophoresis separation, proteins were transferred onto nitrocellulose membranes and blocked with 5% low-fat milk. Membranes were incubated with specific primary antibodies, washed with PBS containing 0.001% Tween-20 (PBST) and incubated with the appropriate horseradish peroxidase-conjugated secondary antibody. After washing in PBST, membranes were subjected to enhanced chemiluminescence detection analysis. For IP analysis, cells were solubilized in lysis buffer (see above). Cell lysates were incubated with the specific antibody for 2–3 h, at 4 °C followed by 3–18 h rotated incubation with protein A/G agarose beads (Santa Cruz Biotechnology, Santa Cruz, CA, USA) at 4 °C. Beads were collected by slow centrifugation, washed four times with lysis buffer and analyzed by SDS–polyacrylamide gel electrophoresis followed by detection with specific antibody. IP from medium was performed similarly.

## Author Contributions

O. Shomron, I. Nevo-Yassaf, T. Aviad, J. Shepchelovich, Y. Yaffe, E. Erez Zahavi, Anna Dukhovny, E. Ella H. Sklan, Perlson and Y. Yonemura carried out experiments.

A. Yeheskel, M. Pasmanik-Chor performed bioinformatics analysis.

G. H. Patterson carried out MSIM analysis.

O. Shomron, C. Kaether, and K. Hirschberg wrote the manuscript Anna Mironov, Galina V Beznoussenko, Alexander A. Mironov carried out electron microscopy experiments

## Acknowledgments

Thanks to Ben Nichols (MRC, United Kingdom), Jennifer Lippincott Schwartz, Janelia, VA.,USA), Nihal Altan Bonnet (NIH, MA, USA) Ella Sklan (TAU, Israel) for critical reading and suggestions, Ron Kopito (Stanford, CA, USA) for the CFTR-GFP constructs. Tom Rapoport (Harvard, Boston) for the reticulon-GFP, Rina Harbesfeld (TAU, Israel) for CPE-GFP. Vivek Malhotra for procollagen-I (CRG, Barcelona, Spain) This work was supported by ISF grant 1063/16 to KH. Thanks to the Imaging facility at FLI, Jena, Germany.

## Supplementary movie caption and figure legend

### Movie caption

**Movie S3A.** Comparison between membrane-associated COPII (red) and cargo protein VSVG-YFP (green) dynamics. Time lapse image sequence captured at 0.66 sec intervals of COS7 cell coexpressing VSVG-YFP (green) and the COPII subunit Sec24C-mCherry (red).

**Movie S3B.** Comparison between membrane-associated COPII (red) and the growing microtubule plus end marker EB3-GFP (green) dynamics. Time lapse image sequence captured at 3.7 sec intervals of COS7 cell coexpressing EB3-GFP (green) and the COPII subunit Sec24C-mCherry (red).

**Movie S4A.** Export of VSVG-YFP from the ER. Accumulation in ERESs and fission of the cargo protein VSVG-YFP (green) in COS7 cells coexpressing Sec24C-mCherry (red). Images were captured at 9.5 sec intervals. See Figure 3A

**Movies S4B-C.** Cargo accumulation in ERES membranes. Cos7 cells were co-transfected with VSVG-YFP (green) and the COPII subunit Sec24C-mCherry (red) Images were captured at 2.6 sec intervals.

**Movie S4D-E.** Fission of transport carriers containing VSVG-YFP (green) from ERESs in COS7 cells co-expressing Sec24C-mCherry (red). Images were captured at 2.6 sec intervals.

**Movie S4F.** Localization of the transport machinery protein Rab1-mCherry (red) in ER to Golgi carriers (arrowheads) containing VSVG-YFP (green) in Huh7 cells.

**Movie S5J.** An ERES connected to the ER via a tubular-shaped membrane in a living cell coexpressing VSVG-YFP (green) and the ER luminal marker signal sequence-mCherry (red) under BFA/Noc-treatment.

**Movie S5Ka-b.** The COPII-bound membranes are localized between ERES and ER. Deconvolved confocal image series from living intact COS7 cells coexpressing Sec24C-mCherry (red) and VSVG-YFP (green) in the presence of BFA/Nocodazole.

**Movie S6A.** Time lapse sequence of COPII-mediated cargo concentration dynamics in ERESs of BFA/Noc-treated COS7 cells. Cells coexpressing VSVG-YFP (green) and Sec24C-mCherry (red) were imaged at 0.2 sec intervals.

**Movie S6B.** Time lapse sequence of cargo and membrane flow into ERESs in BFA/Noc-treated COS7 cells expressing VSVG-YFP.

## Supplementary figures

**Figure S1.**
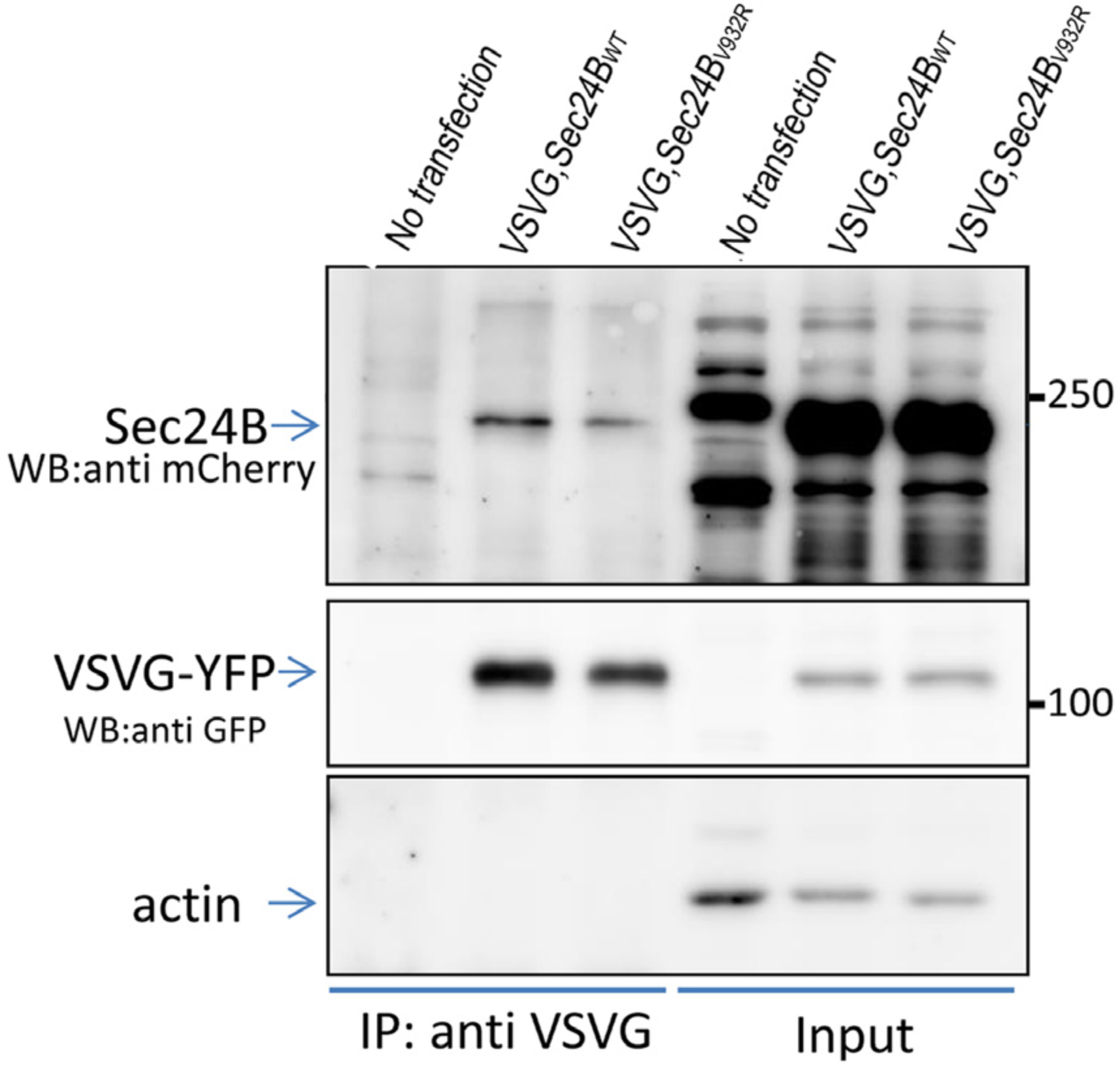
The Sec24B_V932R_ mutant interacts with VSVG. Immunoprecipitation analysis demonstrating interaction between the Sec24B_V932R_ mutant and VSVG. Cells coexpressing VSVG-YFP and wild type or mutant Sec24B were immunoprecipitated with anti-VSVG Abs, separated on SDS-PAGE, blotted and probed with anti-GFP and anti-mCherry for VSVG and Sec24B respectively.

**figure S2.**
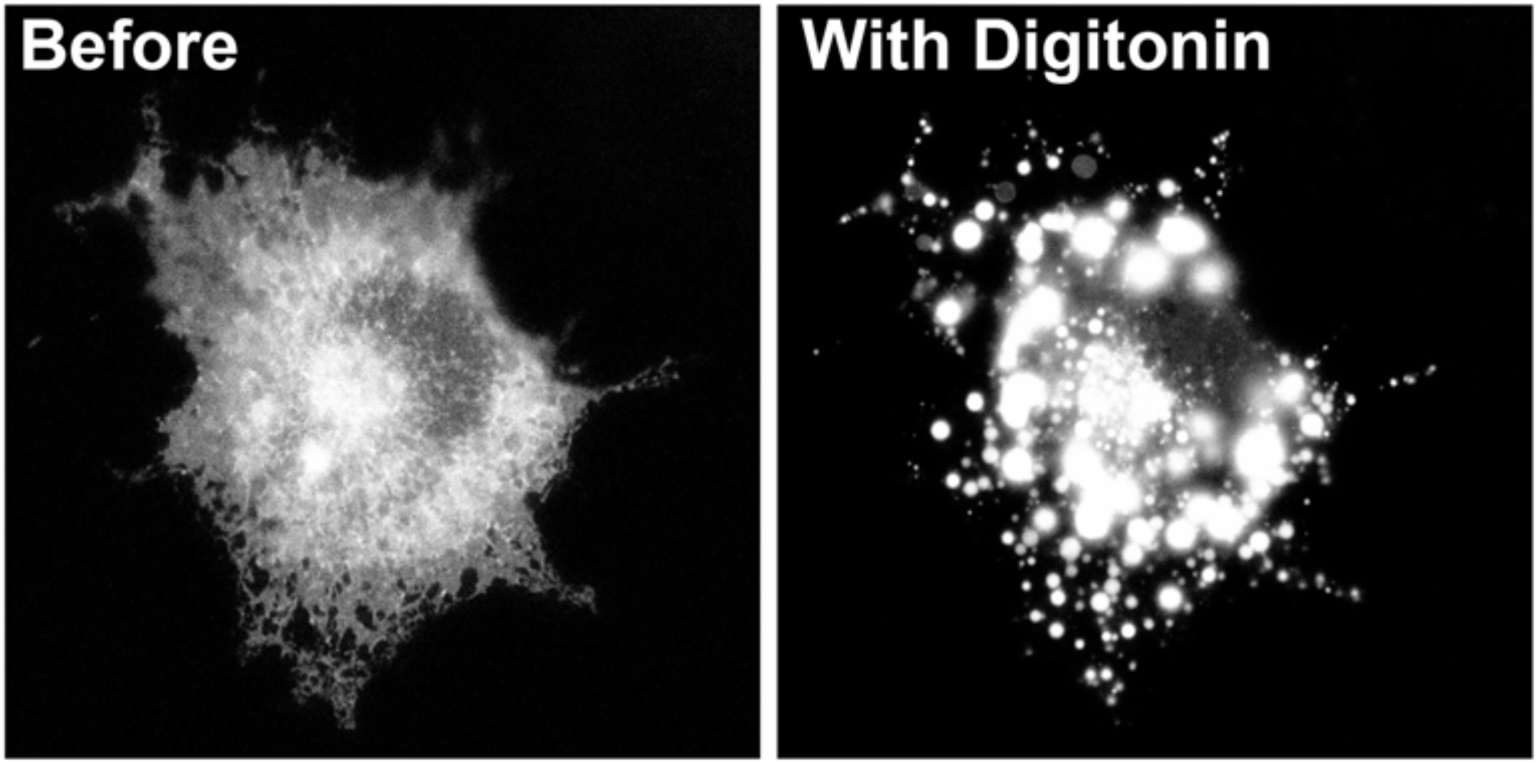
Effect of Digitonin on organelle endomembranes. A living Cos7 cells expressing the ER luminal marker ssmCherry was treated with 40µg/ml digitonin for 1-2 minutes and imaged using …

